# Accelerating antimalarial drug discovery with a new high-throughput screen for fast-killing compounds

**DOI:** 10.1101/2024.04.21.590452

**Authors:** Takaya Sakura, Ryuta Ishii, Eri Yoshida, Kiyoshi Kita, Teruhisa Kato, Daniel Ken Inaoka

## Abstract

The urgent need for rapidly acting compounds in the development of antimalarial drugs underscores the significance of such compounds in overcoming resistance issues and improving patient adherence to antimalarial treatments. The present study introduces a high-throughput screening (HTS) approach using 1536-well plates, employing *Plasmodium falciparum* lactate dehydrogenase (PfLDH) combined with nitroreductase (NTR) and fluorescent probes to evaluate inhibition of the growth of the asexual blood stage of malaria parasites. This method was adapted to efficiently measure the parasite reduction ratio (PRR) in a 384-well plate format, streamlining the traditionally time-consuming screening process. By successfully screening numerous compounds, this approach identified fast-killing hits early in the screening process, addressing challenges associated with artemisinin-based combination therapies. The high-throughput PRR method is expected to be of value in continuously monitoring fast-killing properties during structure-activity relationship studies, expediting the identification and development of novel, rapidly acting antimalarial drugs within phenotypic drug discovery campaigns.

## Introduction

Malaria is a parasitic disease caused by *Plasmodium* spp. infection, and typically is seen in tropical and subtropical areas. In 2021, over 420,000 deaths, predominantly among children under 5 years old, were reported worldwide, along with more than 200 million cases^1^. Artemisinin-based combination therapies (ACTs) are common treatments for malaria in endemic areas. Unfortunately, artemisinin-resistant parasites have spread across Southeast Asia and Africa, highlighting the need for new antimalarial drugs to combat this life-threatening parasitic disease^2–4^. In the last decade, the combination of phenotypic screening, and *in vitro* evolution and whole-genome analysis (IVIEWGA) has played a significant role in antimalarial drug development. This approach has contributed to the identification of several novel drug targets for malarial parasites, facilitating target-based HTS campaigns ^5,6^. Despite the strong efforts in target-based drug discovery being conducted by the Medicines for Malaria Venture (MMV) and collaborating researchers, the development of novel antimalarial medicines often has been hampered by resistance of the malarial parasites to the drug candidates. For example, recrudescent parasites emerged in Phase II clinical trials of both of DSM265 and cipargamin that targets dihydroorotate dehydrogenase (DHODH) and the *Plasmodium falciparum* Na^+^ pump PfATP4, respectively^7–9^. Although target-based drug discovery is an effective method for generating lead compounds, the targets identified by IVIEWGA carry the risk of potential resistance, a major concern in the later stages of drug development. Therefore, there has been renewed interest in phenotypic screening, with several of the resulting candidates now listed in the current drug development pipeline of MMV.

Single-exposure radical cures (SERCs) are highly appealing, since this strategy prevents parasite resistance and simplifies patients’ treatment. However, SERCs characteristically require fast-killing, as seen in next-generation lead compounds such as MMV688533, INE963, ZY19489, and GSK484, the subjects of ongoing clinical trials^10–13^. The standard method employed to determine the speed of killing of parasites, the parasite reduction ratio (PRR), first was described in 2012; since then PRR has been utilized widely to assess the killing properties of candidate compounds^14,15^. The conventional PRR assay has a relatively low throughput due to the complexity of the protocol, requiring at least one month to yield results. This aspect means that one cannot determine whether a given hit compound is a fast killer until the candidate selection step in the drug development pipeline. Despite the crucial importance of the PRR properties of next-generation lead compounds, no scalable assay method for determining the PRR values of multiple compounds has been reported to date.

Phenotypic screening of the asexual blood stage (ABS) of malaria parasites often employs the *P. falciparum* lactate dehydrogenase (PfLDH) assay to quantify parasite growth^16^. Specifically, PfLDH (in the lysate of a parasite culture) is coupled to the *Clostridium kluyveri* diaphorase; the bacterial enzyme catalyzes a redox reaction that converts nitroblue tetrazolium (NBT) to nitroblue formazan (NBF) via 3-acetyl pyridine adenine dinucleotide (APAD^+^), a nicotinamide adenine dinucleotide (NAD^+^) analogue that exhibits cofactor specificity for PfLDH. In the past, GlaxoSmithKline (GSK) employed the PfLDH assay to screen approximately 2 million compounds in a 384-well plate format. However, of late, the SYBR green I-based fluorescence assay, which detects parasite DNA, has become more commonly used for phenotypic HTS of the ABS of malaria parasites. This shift occurred because the fluorescence assay was the only method available for use in the 1536-well format^17,18^. Nevertheless, the simple principle of a PfLDH assay that involves a redox enzyme and a detection probe coupled with the LDH reaction, allows for easy tuning of the assay protocol depending on the specific purpose of the experiment.

In the present report, we describe the development of a modified PfLDH assay using the nitroreducatase (NTR) encoded by the *Escherichia coli nfsB* gene. NTR, a flavoprotein, uses NAD(P)H as an electron donor to reduce a variety of nitroaromatics^19^. To enhance the conventional PfLDH assay’s performance, we incorporated into the assay system “turn-on” fluorescence probes that include a nitrobenzyl moiety^20,21^ that is reduced by NTR. The use of a near-infrared (NIR) probe with an extended wavelength range (excitation/emission wavelengths (λ_ex/em_) = 615/690 nm) compared to haemoglobin absorption (∼600 nm) effectively decreased interference from haemoglobin absorption in the lysate, resulting in an increased signal window. This innovative approach using the NIR probe enabled us to conduct phenotypic screening of the ABS of malaria in a 1536-well format, and improved the assay robustness compared to that of the conventional SYBR green I assay. Moreover, the use of the optimized PfLDH-NTR assay system streamlined the time-consuming protocol of conventional PRR. A feasibility study with representative antimalarial drugs showed results consistent with those obtained using the conventional PRR method. Our high-throughput PRR (HT-PRR) assay therefore is expected to serve as a powerful tool for identifying, in the early stages of drug discovery, novel antimalarial compounds with fast-killing profiles.

## Results

### Coupling of NTR reaction with PfLDH

Histidine-tagged NTR expressed in *E. coli* was purified successfully using conventional nickel-nitrilotriacetic acid (Ni-NTA) agarose purification, yielding a specific activity in the range of 3−5 μmol/min/mg (Fig. 1a). The recombinant NTR converted nitrobenzyl-umbelliferone 1 (NCOU1) to umbelliferone, such that the fluorescent signal of the product (λ_ex/em_ = 315/455 nm) exhibited a time-dependent increase during room temperature incubation (Fig. 1b, c). For our assay, PfLDH was coupled with the purified NTR, rather than with the bacterial diaphorase used in the conventional PfLDH assay, and parasite growth was quantified by monitoring the fluorescence signal of umbelliferone (Fig. 2a). As anticipated, the NTR quantitatively reduced NCOU1 by using the reducing equivalents from the APADH produced by the PfLDH enzyme present in the parasite lysate. A titration of cultured *P. falciparum* using the PfLDH-NTR assay demonstrated that the fluorescent signal increased in a parasitemia-dependent manner, with the highest signal intensity seen under the condition of 1% haematocrit (Fig. 2b and Supplementary Fig. 1).

**Figure 1.**
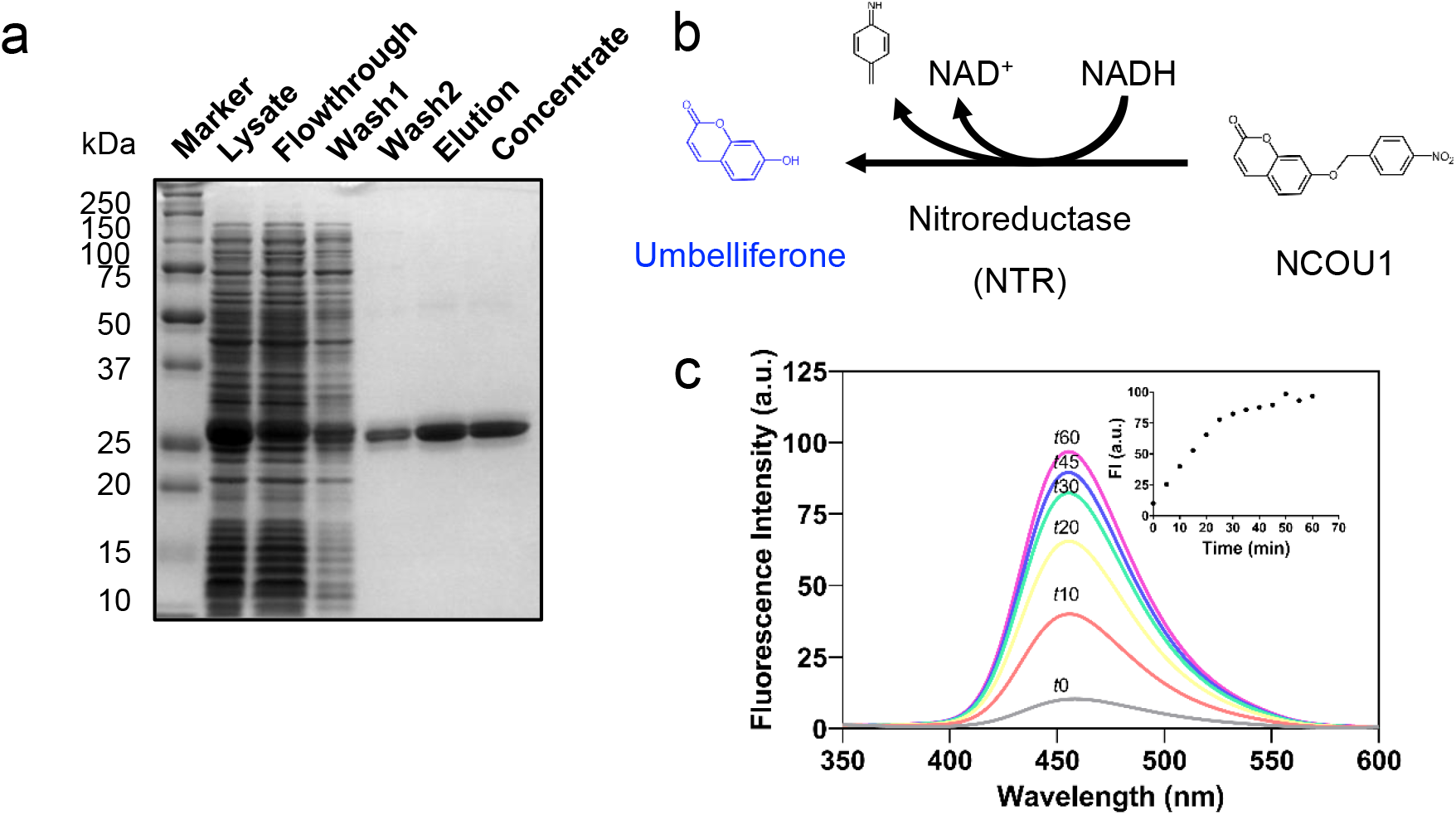
Purification of recombinant nitroreductase (NTR) and quantification of enzymatic activity. **a** Recombinant C-terminally His_10_-tagged NTR was expressed in *E. coli* BL21 Star (DE3) and purified using Ni-NTA agarose. In the lanes denoted as Marker, Lysate, Flowthrough, Wash1 and Wash2, a total of 10 μg protein were loaded. For Elution and Concentrated lanes, 2 μg were loaded. Estimated protein size is 26.7 kDa. **b** Scheme of the enzymatic reaction catalyzed by NTR reducing NCOU1 to umbelliferone (λ_ex/em_ = 315/455 nm). **c** The fluorescent spectra of umbelliferone. Time-dependent increases in the fluorescence spectra were recorded for 60 minutes at room temperature in NTR assay buffer (30 mM Tris-HCl, pH 8.0, 0.25 % Triton X-100) containing 5 μg/mL NTR, 20 μM NCOU1, and 50 μM NADH (λ_ex/em_ = 315/350–600 nm over 0.5-nm intervals, recorded every 5 minutes). For better visualization, only the spectra from time points at 0, 10, 20, 30, 45, and 60 minutes are shown (grey: 0 min, orange: 10 min, yellow: 20 min, light blue: 30 min, blue: 45 min, purple: 60 min). Fluorescence intensity (FI) over time (λ_ex/em_ = 315/455 nm) is plotted in the inset. NCOU1: nitrobenzyl-umbelliferone 1, NAD^+^: oxidized nicotinamide adenine dinucleotide, NADH: reduced nicotinamide adenine dinucleotide, a.u.: arbitrary unit.

**Figure 2.**
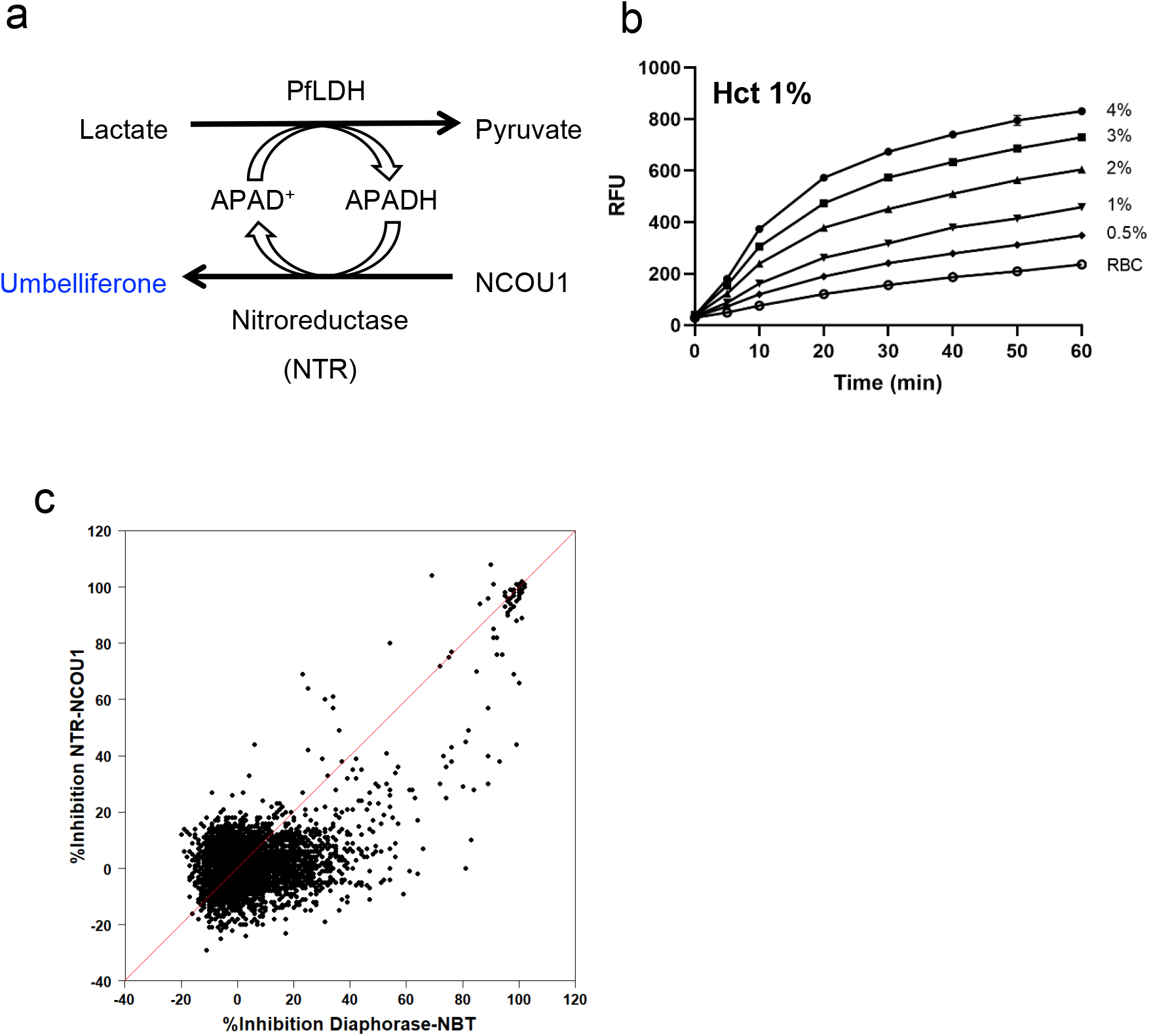
Novel assay using nitroreductase (NTR) coupled with *Plasmodium falciparum* lactate dehydrogenase (PfLDH). **a** Scheme of the PfLDH-NTR coupled reaction used for quantification of parasite growth using NCOU1 as the NTR probe. **b** Parasitemia titration measured by the NTR-NCOU1 assay. The umbelliferone signal increased in a parasitemia-dependent manner. Each parasite culture (1% haematocrit (Hct)) was assayed with PfLDH buffer (100 mM Tris-HCl (pH 8.0), 150 mM lithium L-lactate, and 0.25 % Triton X-100) containing 100 μM APAD^+^, 100 μM NCOU1, and 0.05–0.1 U/mL NTR at room temperature. **c** Correlation of %Inhibition between conventional diaphorase-NBT and NTR-NCOU1 assays of PfLDH activity. The red line indicates unity (y = x). NCOU1: nitrobenzyl-umbelliferone 1, APAD^+^: oxidized 3-acetylpyridine adenine dinucleotide, APADH: reduced 3-acetylpyridine adenine dinucleotide, RFU: relative fluorescence units. RBC: red blood cell.

Next, we compared the screening performance, in a 384-well plate format, of the PfLDH-NTR assay compared to the conventional PfLDH-diaphorase assay. Specifically, we used the two assays to assess the growth inhibition of ABS parasites by a test set of 3,837 compounds from Shionogi & Co., Ltd., using a compound concentration of 2.5 μM. A correlation (r^2^ = 0.387) was observed between the results obtained by the two methods, as depicted in Fig. 2c. While the average signal-to-background (S/B) ratio for the NTR method (5.02) was lower than that of the diaphorase method (8.31), the two assays demonstrated similar Z’-factor values (0.71 and 0.76, respectively) (Table 1). Notably, the diaphorase method produced a higher number of hits (261 hits) than did the NTR method (123 hits). The PfLDH-diaphorase method may have selected for a larger number of pseudo-positive hits, given that this assay’s percent inhibition data did not exhibit a Gaussian distribution and did not correlate with data obtained by other methods (Supplementary Fig. 2, 3, 4). These results indicated that our fluorescence-based PfLDH assay, which incorporates NTR and NCOU1, is more robust than the conventional PfLDH assay using diaphorase.

**Table 1:** Parameters for each assay method.

| Method | Plate<br>format | S/B | CV (%) |  | Z' | Number<br>of hits | Hit rate<br>(%) |
| --- | --- | --- | --- | --- | --- | --- | --- |
|  |  |  | Negative | Positive |  |  |  |
| Diaphorase-NBT | 384 | 8.31 | 8.1 | 2.6 | 0.71 | 261 | 6.8 |
| NTR-NCOU1 | 384 | 5.02 | 5.6 | 3.9 | 0.76 | 123 | 3.2 |
| SYBR green | 1536 | 5.89 | 9.6 | 6.2 | 0.61 | 66 | 1.7 |
| NTR-NIR | 1536 | 11.2 | 8.0 | 6.2 | 0.72 | 70 | 1.8 |

NBT: nitroblue tetrazolium, NTR: nitroreductase, NCOU1: nitrobenzyl-umbelliferone 1, NIR:
near-infrared, S/B: signal/background ratio, CV: coefficient of variation

### NIR probe with PfLDH-NTR assay improves screening robustness

The PfLDH-NTR-NCOU1 assay was not suitable for a 1536-well format due to the low S/B ratio. Consequently, our next objective was to improve this assay method using another probe with a different wavelength of fluorescence emission. Given that the parasite lysate contains a substantial amount of haemoglobin from red blood cells, we conjectured that the absorption of haemoglobin (∼600 nm) interferes with the fluorescent signal of umbelliferone^20^. Therefore, we introduced another NIR probe, one that exhibits longer excitation and emission wavelengths after reduction by NTR (Fig. 3a). We observed that, following reduction of the NIR probe by NTR in the assay solution, the signal intensity of the reduced species (NIR-P^red^), increased in a time-dependent manner, permitting quantification of the malaria parasites (Fig. 3b, c). Given the read-time need for measurement of output in the 1536-well format, we quenched the PfLDH reaction by adding a high concentration of pyruvate (0.9 M), resulting in the complete inhibition of NCOU1 reduction (Fig 3c). For reactions run in the 1536-well format, the parasite growth inhibition detected by the PfLDH-NTR-NIR assay showed a strong correlation with that obtained by the SYBR green I-based assay (Fig. 3d). Notably, the PfLDH-NTR-NIR assay achieved a higher S/B ratio and Z’-factor (11.2 and 0.72, respectively) than those measured by the SYBR green I assay (5.89 and 0.61), indicating that PfLDH-NTR-NIR assay is more robust than the existing method when employed in a 1536-well format.

**Figure 3.**
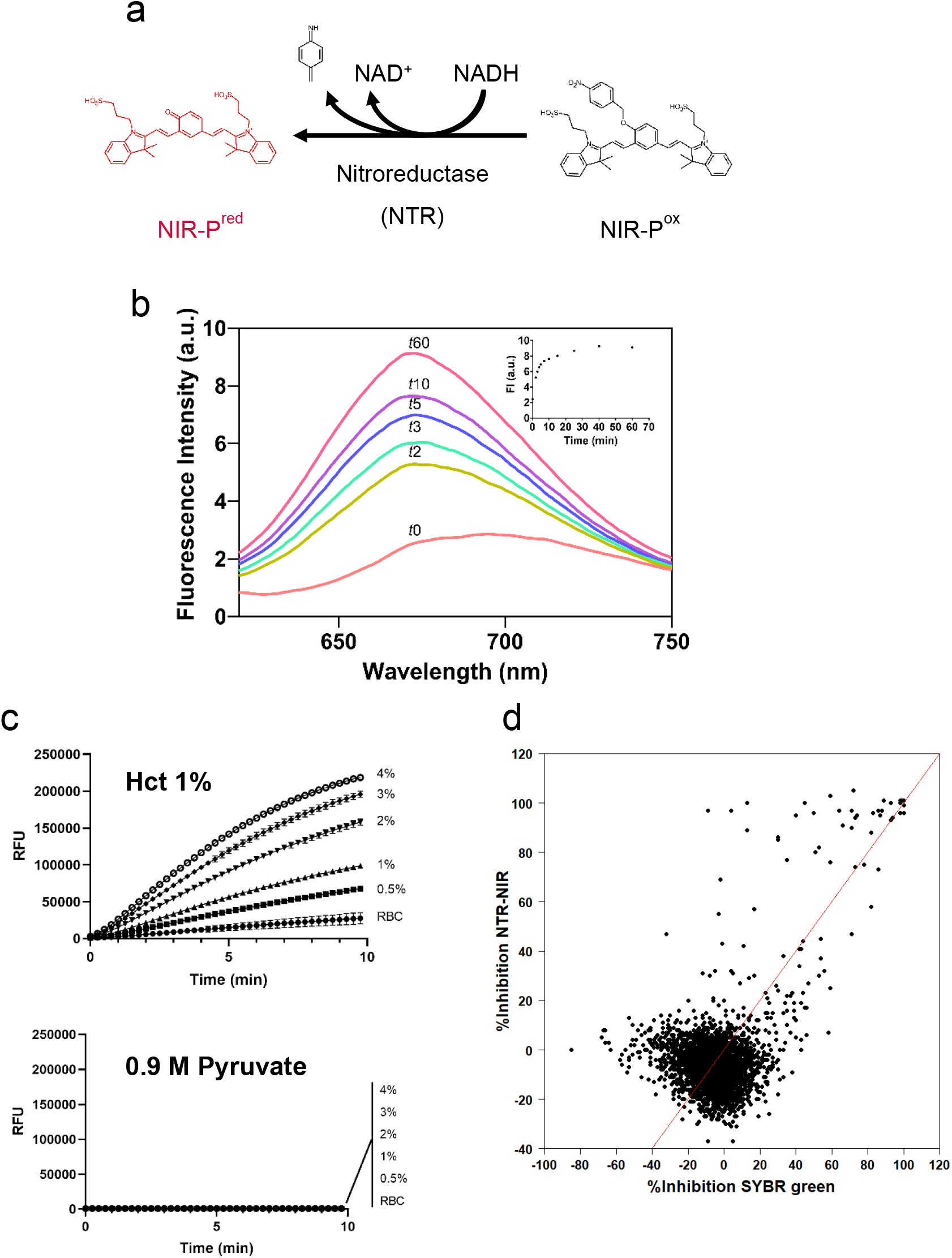
Nitroreductase (NTR) assay using near-infrared probe (NIR-P). **a** Scheme of enzymatic reaction of NTR and NIR-P^ox^ producing the fluorescent NIR-P^red^ (λ_ex/em_ = 615/670 nm). **b** Time-dependent fluorescent spectra of NIR-P^red^ produced by NTR. The enzymatic reaction catalyzed by NTR was monitored using 50 μM NIR-P^ox^ and 50 μM NADH in 30 mM Tris-HCl, pH 8.0, buffer containing 0.5 μg/mL NTR. Fluorescence signals (λ_ex/em_ = 605/620–740 nm over 1-nm intervals) were measured over a 60-minute period at room temperature (orange: 0 min, yellow: 2 min, light green: 3 min, blue: 5 min, purple: 10 min, red: 60 min). Fluorescence intensity (FI) over time is plotted in the inset. Note that the reduction of NIR-P by NTR is much faster than the reaction observed using the NCOU1 substrate (Fig. 1c). **c** Parasitemia titration measured using the NTR-NIR assay. Each parasite culture (1% haematocrit (Hct)) was assayed in PfLDH buffer (100 mM Tris-HCl, pH8.0, 300 mM lithium L-lactate, and 0.25% Triton X-100) containing 500 μM APAD^+^, 200 μM NIR probe, and 0.05 U/mL NTR at room temperature. The quenching (“stop”) solution consisted of 0.9 M pyruvate. Data are presented as the mean ± standard deviation (SD) (n=4). **d** Correlation of %Inhibition between conventional SYBR green and NTR-NIR assays of PfLDH activity. The red line in the graph represents the line of equality (y = x). a.u.: arbitrary unit, NAD^+^: oxidized nicotinamide adenine dinucleotide, NADH: reduced nicotinamide adenine dinucleotide, RFU: relative fluorescence units.

### High-throughput PRR (HT-PRR) assay

In the ABS, LDH oxidizes NADH and provides NAD^+^ to maintain the flux of anaerobic glycolysis required for the synthesis of ATP, a function that is essential for parasite survival^22,23^. We conceived the idea of using PfLDH activity as an indicator of parasite viability and applied the PfLDH-NTR-NCOU1 method in a 384-well format to quantify the PRRs of test compounds. Mixed-stage (non-synchronized) parasites at 1.5% parasitemia were incubated at 37 °C for 0, 24, 48, 72, and 96 hours with test compounds at concentrations of 10 μM. Live parasite signals were quantified by the PfLDH-NTR-NCOU1 method and the PRR values were calculated for each compound. The values then were normalized (as a percentage) to those of dihydroartemisinin (DHA; defined as 100%), a representative fast-killing compound, and ELQ300 (defined as 0%), an inhibitor of the electron transport chain (ETC) that is classified as a slow-killing compound (Supplementary Fig. 5). At 24 hours, fast-killing compounds like ACT-451840, artemether, and cipargamin exhibited PRR values exceeding 70%; chloroquine, MMV048, and ganaplacide, categorized as moderate-speed-killing compounds, yielded PRR values in the approximate range of 40–70%. Other known slow-killing ETC inhibitors, DSM265 and atovaquone, yielded lower PRR values (16.9 and 8.6%, respectively). These results are consistent with the data obtained by the conventional PRR method. The sole exception was DDD107498, a compound known to be a slow-killing inhibitor by conventional PRR^24^, show high PRR value. Given that this compound is an inhibitor of the eukaryotic translation elongation factor 2 (eEF2)^24^; we hypothesize that the interruption of protein synthesis by DDD107498 may directly impact PfLDH synthesis, leading to lowered enzymatic activity, which our assay detects as an elevated PRR.

Subsequently, we sought to optimize the level of parasitemia in the assay by employing our method to assess the effects, on parasites at various densities, of 24- and 48-hour exposure to the known inhibitors atovaquone, DHA, and cipargamin (Supplementary Fig. 6). We observed that when the starting parasitemia exceeded 2%, the signal for vehicle (dimethylsulfoxide; DMSO)-treated parasites achieved a virtual plateau after 48 hours of incubation. Therefore, we fixed the starting parasitemia at 1.5% for further experiments.

Having completed the optimization of HT-PRR, we conducted PRR assays in an 8-point dose-response experiment. The dose-response PRR experiment was implemented because during the test run, we observed that the shape of PRR dose curves differed among the compounds (Supplementary Fig. 7). The experiment assessed PRR at intervals of 0, 24, 48, and 72 hours; and determined the 50% effective concentration to reduce the PRR values (PRR_50_) of the representative test compounds, in addition to the 72-hour cultures for standard growth inhibition (EC_50_) assays (Fig. 4a). A comparison of the inhibition curves revealed that Hill slopes and plateau levels of the PRR curves varied among compounds and time points. To quantify the shape of the dose-response curve and utilize these data as an indicator of PRR, we calculated for each compound the ratio of area under the curve (AUC) between the growth inhibition curve and the PRR curve, yielding a parameter that we refer to as the “AUC%” (Table 2). We observed that the AUC% values of the fast-killing compounds exceeded 80%. In contrast, the slow-killing compound exhibited lower AUC% values (of approximately 50%); furthermore, the % inhibition by slow-killing compounds did not reach 100%, even at the highest tested concentration (1 μM). These data suggested that robust AUC% values, calculated using multiple data points, permitted effective classification of compounds according to their killing-rate properties.

**Figure 4.**
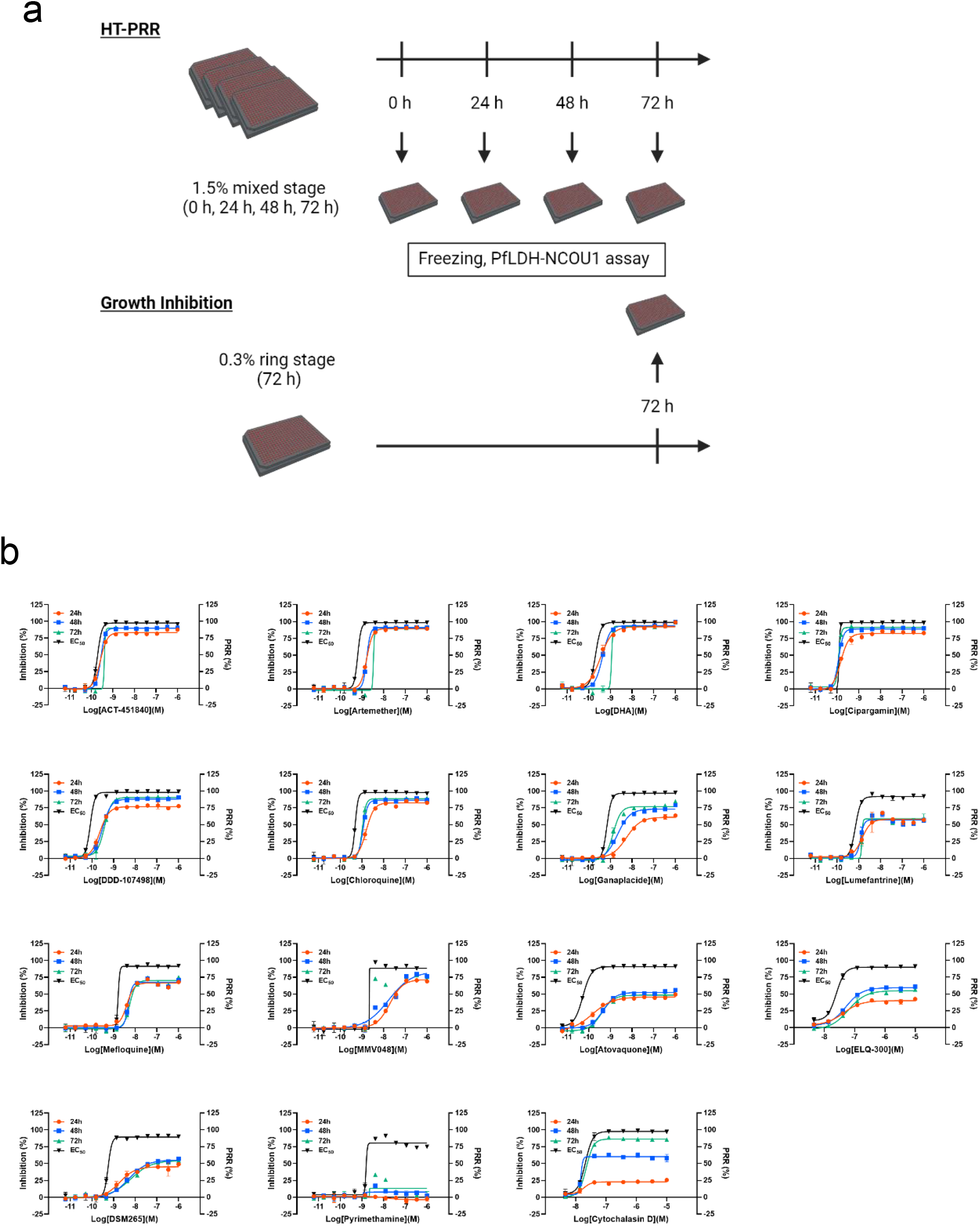
Schematic representation of high-throughput parasite reduction ratio (HT-PRR) and assessment of well-characterized antimalarial agents. **a** To evaluate the parasite reduction ratio (PRR) by established HT-PRR methods, non-synchronized 3D7 parasites (haematocrit (Hct) 1%, parasitemia 1.5%) were plate-cultured for 0, 24, 48, and 72 hours in the presence of compound. For the growth inhibition assay, 3D7 parasites were synchronized by exposure to 5% sorbitol; ring stage parasites then were plate-cultured for 72 hours in the presence of compound. Growth was terminated at the indicated time point by freezing (at -30 ° C) the respective assay plates; the number of live parasites then was quantified via an LDH assay using retororeductase (NTR) and nitrobenzyl-umbelliferone 1 (NCOU1). For the growth inhibition assay, the positive control consisted of a mixture of 1 μM atovaquone and artemisinin; the negative control consisted of 0.4% dimethylsulfoxide (DMSO). For the HT-PRR assay, the positive and negative controls consisted of 1 μM dihydroartemisinin (DHA) and 0.4% DMSO, respectively. **b** Dose-response curves of Inhibition (%) and PRR (%) by test compounds were generated for each assay condition (n =2).

**Table 2:** Activity and AUC% of test compounds.

| Compound | PRR<sub>50</sub> (nM) |  |  | EC<sub>50</sub> (nM) | AUC% |  |  |
| --- | --- | --- | --- | --- | --- | --- | --- |
|  | 24 h | 48 h | 72 h |  | 24 h | 48 h | 72 h |
| ACT-451840 | 2.77 | 2.72 | 4.07 | 1.95 | 89.2 | 95.6 | 90.6 |
| Artemether | 16.5 | 16.4 | 35.2 | 6.34 | 87.1 | 88.2 | 77.3 |
| DHA | 3.36 | 4.13 | 11.8 | 2.05 | 94.2 | 93.9 | 81.8 |
| Cipargamin | 1.61 | 1.17 | 1.15 | 1.29 | 88.6 | 98.2 | 100.1 |
| DDD-107498 | 2.70 | 3.12 | 3.67 | 0.84 | 76.1 | 84.0 | 85.0 |
| Chloroquine | 16.9 | 11.3 | 10.4 | 4.85 | 78.6 | 86.6 | 89.1 |
| Ganaplacide | 117.5 | 30.0 | 15.9 | 7.05 | 55.1 | 71.9 | 80.0 |
| Lumefantorine | 24.7 | 15.3 | 15.0 | 7.28 | 67.9 | 67.4 | 66.7 |
| Mefloquine | 52.9 | 54.5 | 66.7 | 15.9 | 76.1 | 72.3 | 71.7 |
| MMV048 | 276.9 | 209.6 | 23.5 | 23.9 | 62.6 | 72.1 | 78.3 |
| Atovaquone | 124.9 | 15.0 | 20.8 | 0.57 | 49.5 | 51.4 | 46.9 |
| DSM265 | 1186 | 211.5 | 332.4 | 5.84 | 50.6 | 51.7 | 50.2 |
| ELQ-300 | 9990 | 139.3 | 298.5 | 24.8 | 40.3 | 56.9 | 48.5 |
| Pyrimethamine | 9990 | 9990 | 9990 | 15.3 | 10.1 | 21.5 | 0.00 |
| Cytochalasin D | 9990 | 15.9 | 24.1 | 20.5 | 24.7 | 65.3 | 86.8 |

PRR<sub>50</sub>: Concentration of the compounds required to reduce the PRR values to 50%; EC<sub>50</sub>:
effective concentration to reduce the parasite growth to 50%; AUC: area under the curve; DHA:
dihydroartemisinin

## Discussion

In the present work, we established a novel PfLDH-based assay method for detecting antimalarial candidates targeting the ABS by using NTR and “turn-on” fluorescent probes with a nitrobenzyl moiety. The new method provides an assay with robustness superior to that of existing screening methods, and we successfully conducted HTS with a small compound library in a 1536-well format. Our method also was able to assess the parasite survival rate, as assessed by PfLDH activity. The PfLDH-NTR assay permits quantification of the PRR for many compounds in a 384-well format, significantly decreasing experimental time. The PRR classification of known antimalarial drugs by our HT-PRR method was consistent with the data obtained by conventional assays. Moreover, HT-PRR permitted the determination of the dose-dependency of a compound’s PRR, yielding a new parameter (the AUC%) indicating the speed of killing. Our novel PfLDH-NTR-based assay is expected to be valuable for determining the PRR values of many hit compounds in the early stages of drug discovery, facilitating the development of next-generation fast-killing antimalarial drugs.

In the field of cancer research, NTR is widely used for the *in vivo* detection of tumour cells and the activation of prodrugs, reflecting the fact that NTR expression is specifically increased in tumour cells growing under hypoxic conditions^25–27^. Additionally, NTR plays a crucial role in the treatment of Chagas disease, a condition caused by *Trypanosoma cruzi* infection; benznidazole and nifurtimox, front-line drugs used against Chagas, are activated by NTR activity. Resistance to these drugs is determined primarily by loss of function mutations in the *T. cruzi* NTR-encoding gene (including homozygous nonsense mutations and the loss of one copy) that lead to decreased levels of the trypanosomal enzyme^28,29^. Given NTR’s biological significance and diverse applications in biomedical research, we envisioned using NTR to establish innovative assays for the discovery of antimalarial drugs. Moreover, the tissue transparency of the λ_ex/em_ of NIR, which fall within the bio-optical window (650–1100 nm), makes these wavelengths useful in biomedical research, especially for *in vivo* imaging, due to lower haemoglobin absorption^30^. This property of the NIR probe fluorophore mitigates interference from the abundant haemoglobin in red blood cells, greatly enhancing the signal strength of the PfLDH assay. Through the introduction of a coupling enzyme and probes, we improved the performance of the conventional PfLDH assay for detecting growth inhibition of the malaria parasite at the ABS. In the future, it will be interesting to introduce modifications to existing assay systems, thereby enhancing robustness and reducing cost. This initiative has the potential to pave the way for the creation of advanced HTS systems applicable to other infectious diseases, including *Trypanosoma* spp. and *Mycobacteria* spp.; indeed, revisions to facilitate screens for these systems are currently in progress.

For HTS of the ABS of the malaria parasite in a 1536-well format, the commonly used assay employs SYBR green I, a fluorescent dye with excitation and emission wavelengths of 485 nm and 528 nm, respectively. Our PfLDH-NTR assay with NIR^ox^ would be (to our knowledge) the first system using NIR wavelengths to screen for antimalarial compounds. A noteworthy finding is that a small number of hit compounds of the SYBR green I-based assay was not common with that of the PfLDH-NTR with NIR^ox^ assay (Supplementary Fig. 4). The optical properties of small organic compounds in chemical libraries may impact the readout signal, resulting in false-positives and -negatives^31,32^. Indeed, a small number of compounds showed less than 40% inhibition as assessed by the SYBR green I-based assay, possibly reflecting the optical properties of these compounds (Supplementary Fig. 2). The PfLDH-NTR with NIR^ox^ assay may serve as a viable alternative screening method to prevent the loss of promising compounds as pseudo-negatives in future phenotypic HTS of the malaria parasite at the ABS.

To enhance the throughput in assessing PRR, we developed the PfLDH-NTR assay system, enabling the HTS evaluation of antimalarial compounds based on PRR values. Notably, we observed that the dose dependent PRRs of fast-killing compounds typically demonstrate curves that resembled the growth inhibition curve of the same chemicals. However, disparities between these two types of curves are more pronounced for moderate- and slow-killing compounds. To quantify these differences, and to classify multiple test compounds according to the PRR value, we defined the AUC%. This parameter provides a succinct description of the parasite killing rate of test compounds and is expected to accelerate structure-activity relationship (SAR) studies, particularly for fast-killing compounds. At the same time, we note that a limitation of HT-PRR is that PRR values for compounds targeting protein synthesis (e.g., DDD107498) may be overestimated due to the dependence of the assay on PfLDH activity. However, this limitation can be addressed by cross-referencing results with conventional PRR or classical Giemsa staining at later stages of drug discovery.

## Conclusion

We developed, in-house, NTR-based methods for the identification of compounds with activity against the ABS of malaria parasites, and successfully implemented one such assay for HTS in a 1536-well format. The use of the PfLDH-NTR method for PRR determination represents a novel assay of the parasite-killing rate of drug candidates. These innovative new HTS methods are expected to permit the identification of fast-killing compounds with novel scaffolds and properties, paving the way for the discovery of next-generation antimalarial drugs that may find use in combatting drug resistance in malaria-endemic areas.

## Methods

### Preparation of recombinant enzymes

Prior to assay development, we prepared two coupling enzymes, *Clostridium kluyveri* diaphorase and *Escherichia coli* NAD(P)H nitroreductase NfsB (NTR), as previously described^19,33^. Briefly, codon-optimized genes were synthesized to encode proteins with N-terminal His10-SUMO tags and C-terminal His10 tags. These genes with C-terminal His10 were cloned separately into the pET101-D-TOPO expression vector (Thermo Fisher Scientific), and the resulting plasmids were transformed into *E. coli* BL21^TM^ Star (DE3) cells according to the manufacturer’s protocol. The resulting *E. coli* strains were pre-cultured (overnight at 37 °C with shaking at 200 rpm) in Luria– Bertani medium supplemented with 100 μg/mL carbenicillin. The resulting cultures were used as inocula for 600-mL volumes of Terrific-Broth medium supplemented with 100 μg/mL carbenicillin and 0.4% glycerol, and the resulting cultures were incubated at 37 °C with shaking at 200 rpm. When the optical density at 600 nm (OD_600_) reached 0.4 to 0.6, protein expression was induced by adding 50 μM isopropyl β-D-1-thiogalactopyranoside (IPTG; Sigma) and 1 mg/mL riboflavin (Sigma). After incubation for 16 h at 20 °C, the cells were harvested by centrifugation at 7,000 × *g* for 10 min at 4 °C and re-suspended at a density of 0.4 g cell pellet/mL in cold lysis buffer (50 mM potassium phosphate buffer, pH 8.0, 300 mM NaCl, 0.5 mM ethylenediaminetetraacetic acid (EDTA) and 0.25 mM phenylmethylsulfonyl fluoride (PMSF)). The suspended cells were lysed using a French press (Ohtake) at 180 MPa, and the supernatant was collected by centrifugation at 40,000 × *g* for 30 min at 4 °C. The His-tagged proteins were purified using Ni-NTA (Qiagen) following the manufacturer’s protocol and eluted using lysis buffer containing 200 mM imidazole. The eluate was concentrated using an Amicon Ultra Centrifugal Filter 10 kDa Molecular Weight Cut-off (Merck) according to the manufacturer’s protocol, and the resulting purified enzyme was mixed with an equal volume of cold glycerol and stored frozen at -30 °C. The protein concentrations in the final enzyme stocks were determined by the Bradford assay using a Bio-Rad kit, and the specific activities of the enzymes were quantified by measuring the reduction of 2,6-dichlorophenolindophenol (DCIP; Sigma) using a UV760 spectrophotometer (Jasco). To confirm the purity of the enzymes, proteins were separated by sodium dodecyl sulfate-polyacrylamide gel electrophoreses (SDS-PAGE) and stained using GelCode™ Blue Safe Protein Stain (Thermo Fisher Scientific). The specific activities of diaphorase and NTR were typically in the range of 100–150 and 3–5 μmol/min/mg, respectively.

### Quantification of the reduction of fluorescence probes by NTR

To test whether the purified NTR was capable of reducing fluorescence probes, we initially measured NTR activity prior to using the enzyme for parasite quantification. Using a cuvette, 20 μM NCOU1 (Enamine) was combined with a mixture containing 5 μg/mL NTR, 50 μM NADH, and 0.25 % Triton X-100 in 30 mM Tris-HCl (pH 8.0), and the fluorescence of the umbelliferone (λ_ex_/_em_ = 315/455 nm) generated by the reaction was measured. In addition, we synthesized the NIR^ox^ probe according to the previously described route^21^. For the NIR reaction, 50 μM NIR-P^ox^ was combined with a mixture containing 0.5 μg/mL NTR and 50 μM NADH in 30 mM Tris-HCl (pH 8.0), and the fluorescence of NIR-P^red^ (λ_ex_/_em_ = 615/670 nm) was measured. The fluorescence kinetics of both reduced probes were monitored using a FP-6300 spectrofluorometer (Jasco).

### Parasite cultures

*Plasmodium falciparum* 3D7 parasites were maintained in RPMI1640 medium supplemented with 25 mg/L gentamicin, 50 mg/L hypoxanthine, 23.8 mM sodium bicarbonate, and 0.5% (w/v) Albumax II (Gibco). The culture was maintained at 2% haematocrit using O^+^ human erythrocytes obtained from The Japanese Red Cross Society. As per established protocols^34,35^, the incubation temperature was set at 37 °C and the environment consisted of mixed gas (5% O_2_, 5% CO_2_, 90% N_2_). Parasitemia was determined using either Giemsa staining or an XN-30 automated haematology analyser (Sysmex)^36^. For plate assays, ring-stage synchronized parasites were prepared using sorbitol treatment^37^. In the growth inhibition assay, the compound solvent DMSO and a mixture of 1 μM artemisinin/atovaquone served as negative and positive controls, respectively.

### Diaphorase-NBT and NTR-NCOU1 assay in 384-well plates

Test compounds were pre-dispensed at a volume of 100 nL/well into 384-well plates. For the diaphorase-NBT assay, ring-stage parasites at 0.3% parasitemia and 2% haematocrit were dispensed into 384-well clear plates (Corning); for the NTR-NCOU1 assay, ring-stage parasites at 0.3% parasitemia and 1% haematocrit were dispensed into 384-well black plates (Greiner). Parasites then were dispensed at 25 μL/well into the compound-containing plates using a Multidrop Combi with a small metal cassette (Thermo Fisher Scientific). The parasite cultures in the assay plates then were incubated in a moist chamber filled with mixed gas for 72 hours and subsequently frozen overnight (or longer) at -30 °C. The resulting frozen plates were thawed at room temperature for at least 2 hours before the assay was performed. For the diaphorase-NBT assay, the reaction mix was prepared in modified PfLDH buffer (100 mM Tris-HCl, pH 8.0, 150 mM lithium L-lactate, and 0.25% Triton X-100) supplemented with 75 μM APAD^+^, 0.2 mg/mL NBT, and 1 U/mL diaphorase (one unit was defined as the amount of enzyme required to reduce 1 μmol of APADH per minute). For this assay, the reaction mixture was dispensed into the assay plates at 70 μL/well, and the plates then were incubated at room temperature with agitation at 650 rpm for 20 minutes. On the other hand, the reaction mixture for the NTR-NCOU1 assay consisted of modified PfLDH buffer supplemented with 100 μM APAD^+^, 100 μM NCOU1, 0.05 U/mL NTR, and 0.015% (v/v) KM-70 defoaming agent (Shin-Etsu). For this assay, the reaction mix was dispensed into the assay plates at 35 μL/well, and the plates then were incubated under the same condition as those used for the diaphorase-NBT assay. The absorption of nitro blue formazan (650 nm; for the diaphorase-NBT assay) or the fluorescence of umbelliferone (λ_ex/em_ = 360/465 nm; for the NTR-NCOU1 assay) was measured using a SpectraMax® Paradigm spectrophotometer (Molecular Devices, Inc.).

### NTR-NIR^ox^ assay in 1536-well plates

For the HTS in 1536-well format, ring-stage parasites at 0.3% parasitemia and 1% haematocrit were dispensed at 4 μL/well into 1536-well black plates (Greiner) containing 20 nL/well of test compounds. Parasite-dispensed assay plates were maintained under the same conditions as described above for the 384-well plate assays. The NIR^ox^ probe was initially suspended at 1 mM in 90% acetonitrile. The reaction mixture for the NTR-NIR^ox^ assay was prepared by supplementing modified PfLDH buffer with 500 μM APAD^+^, 200 μM NIR^ox^ probe, and 0.05 U/mL NTR. For this assay, the reaction mixture was dispensed into the assay plates at 3 μL/well, and the contents of each well were mixed at 1,200 rpm. The plates then were incubated at room temperature for 3 minutes. The enzymatic reactions were quenched by dispensing the “stop” mixture (0.9 M sodium pyruvate containing 0.015% KM-70) at 3 μL/well, and the assay plates then were centrifuged at 90 × *g* for 1 min at room temperature. The centrifuged plates were incubated for 1 hour at room temperature in a moist chamber, at which point the NIR^red^ signal (λ_ex/em_ = 615/690 nm) was measured using a PHERAstar Plus microplate reader (BMG LABTECH).

### SYBR green assay in 1536-well plates

For the SYBR green assay, ring-stage parasites at 0.3% parasitemia and 3% haematocrit were dispensed at 4 μL/well into 1536-well black plates (Greiner) containing 20 nL/well of test compounds. Parasite-dispensed assay plates were maintained under the same conditions as described above for the 384-well plate assays. The reaction mixture for the SYBR green I assay consisted of lysis buffer (20 mM Tris-HCl, pH 8.0, 5 mM EDTA, and 0.1% Triton X-100) supplemented with 0.02% SYBR green I and 0.015% KM-70. This reaction mixture was dispensed into the assay plates at 4 μL/well, and the contents of each well were mixed at 1,200 rpm. The plates then were incubated at room temperature for 1 hour. The fluorescence of SYBR green I (λ_ex/em_ = 485/528 nm) was measured using a SpectraMax® Paradigm spectrophotometer.

### Data analysis

Hits of a test set of compounds were chosen by activity criteria [>3 standard deviation (SD) cutoff of DMSO controls] that were calculated using Spotfire software (version 11.4.3; TIBCO). Z’ was calculated as 1 − (3 × SD_100%_ + 3 × SD_0%_)/(Mean_100%_ − Mean_0%)_. The correlations of %Inhibition and histograms were visualized with R (version 4.0.2). The inhibition curves and EC_50_ values were determined using GraphPad Prism (version 8.4.3. GraphPad Software, Inc., San Diego, California, USA) and Spotfire. Data visualization for the other figures was performed using GraphPad Prism. The area under the curve (AUC) for both the HT-PRR assay and growth inhibition assay in the 384-well plates was calculated using Spotfire software, and the AUC% was calculated as 100 × [(AUC of HT-PRR assay) / (AUC of NTR-NCOU1 assay)]. Venny (version 2.1) was used to make a Venn diagram.

### HT-PRR assay in 384-well plates

Test compounds were pre-dispensed at 100 nL/well in 384-well plates, and replicate plates were prepared. Non-synchronized parasites at 1.5% parasitemia and 1% haematocrit were dispensed into five 384-well black compound-containing plates at 25 μL/well using a Multidrop Combi with a small metal cassette. One of the plates was frozen at -30 °C right immediately following dispensing of parasites (control plate; 0 hour), while the other plates were incubated in a moist chamber filled with mixed gas for 24, 48, 72, or 96 hours; at the indicated time points, growth was terminated by freezing overnight (or longer) at -30 °C (PRR plates). Additional plates for the generation of the growth inhibition curve were prepared and subjected to the NTR-NCOU1 assay using the procedure described in the previous section, with a 72-hour incubation period. The fluorescence of umbelliferone (λ_ex/em_ = 360/465 nm) was measured using a PHERAstar microplate reader, and the relative LDH activity was calculated by dividing the fluorescence of the PRR plates by that of the respective control plate. The PRR inhibition percentage was calculated by normalizing the primary values to the LDH activity obtained in the negative and positive control reactions (compound solvent DMSO and 1 μM dihydroartemisinin, respectively).

In addition, the concentration of each compound to reduce the PRR value to 50% (PRR_50_) was calculated in each time point.

## Acknowledgements

The authors thank the Japanese Red Cross Society for providing human red blood cells (RBCs; Registry No. R030038) and Dr. Kenji Takaya (Laboratory for Medicinal Chemistry Research, Shionogi & Co., Ltd.) for synthesizing the NIR probe.

## Abbreviations Used

PfLDH: *Plasmodium falciparum* lactate dehydrogenase
NTR: nitroreductase
PRR: parasite reduction ratio
ACTs: artemisinin-based combination therapies
IVIEWGA: *in vitro* evolution and whole-genome analysis
MMV: Medicines for Malaria Venture
DHODH: dihydroorotate dehydrogenase
SERCs: single-exposure radical cures
ABS: asexual blood stage
NBT: nitroblue tetrazolium
NBF: nitroblue formazan
APAD: 3-acetyl pyridine adenine dinucleotide
NIR: near-infrared
NCOU1: nitrobenzyl-umbelliferone 1
ETC: electron transport chain
DHA: dihydroartemisinin
eEF2: elongation factor 2
PMSF: phenylmethylsulfonyl fluoride
DCIP: 2,6-dichlorophenolindophenol

## Author contributions

D.K.I., T.K., and K.K. directed the work. T.S., R.I., and E.Y. performed the experiments. T.S., R.I., and E.Y. analysed the data. T.S., R.I., E.Y., and D.K.I. wrote the paper. All authors directly participated in this work, including preparation of the manuscript and approval of the final version. All authors have read and agreed to the published version of the manuscript.

