## Supplementary figures for "Accelerating antimalarial drug discovery with a new high-throughput screen for fast-killing compounds"

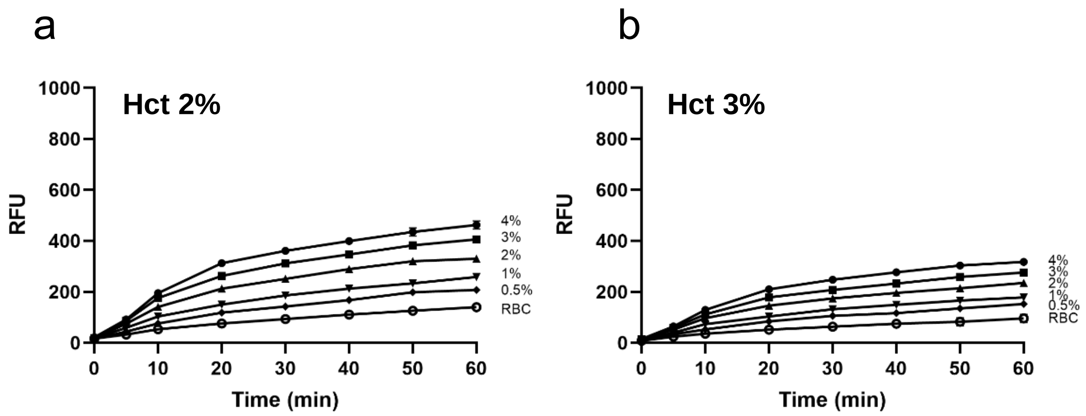


Supplementary Figure 1. Parasitemia titration measured by NTR-NCOU1 assay at 2% haematocrit (Hct) (a) and 3% Hct (b). For better comparison between the datasets, the y-axis scale was designed to match that of the plots in Figures 2b**.** RFU: relative fluorescence units. RBC: red blood cell.

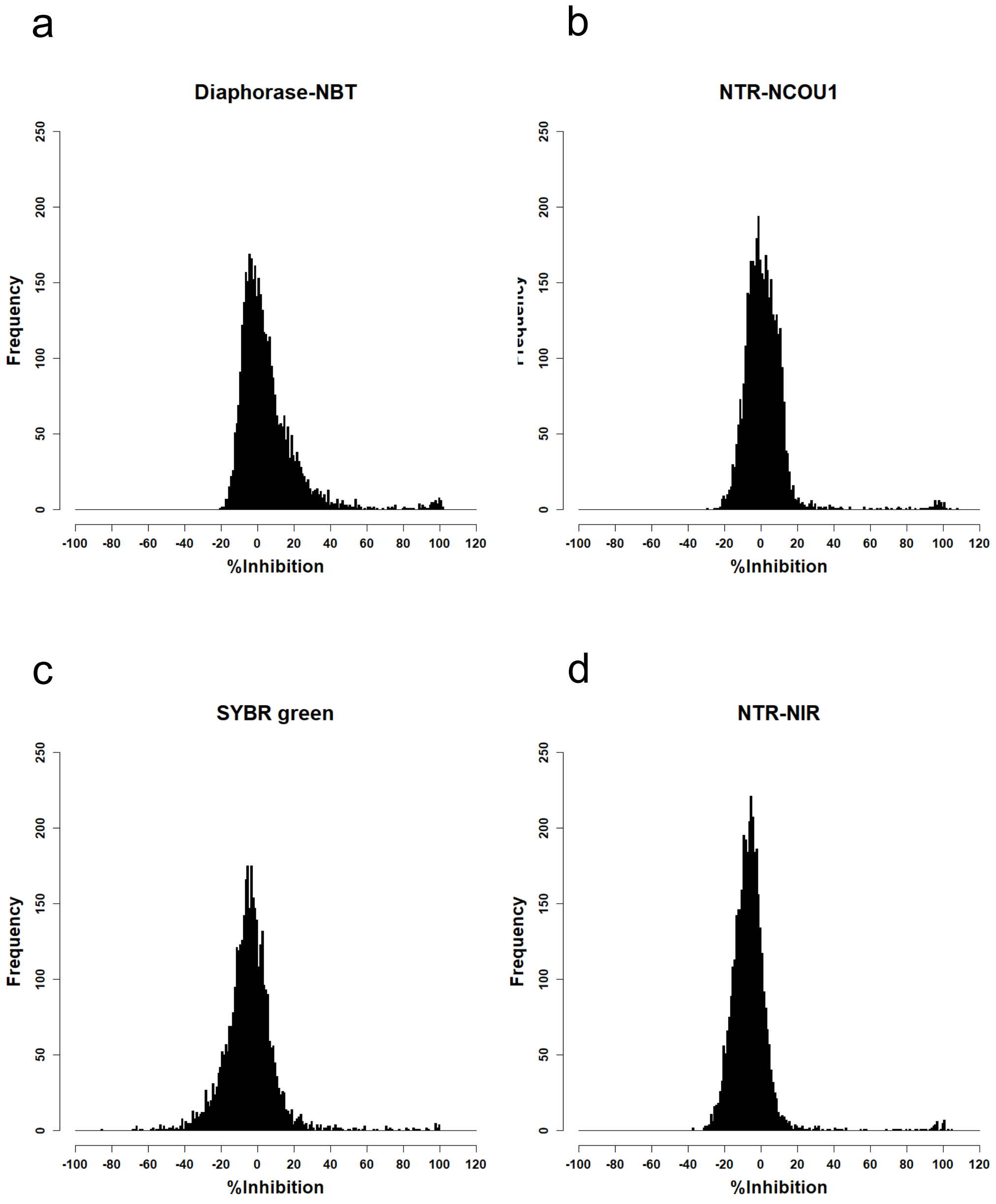


Supplementary Figure 2. Histograms of the hits obtained from using each assay method; Diaphorase-NBT (a), NTR-NCOU1 (b), SYBR green (c), NTR-NIR (d). NBT: nitroblue tetrazolium, NTR: nitroreductase, NCOU1: nitrobenzyl-umbelliferone 1, NIR: near-infrared.

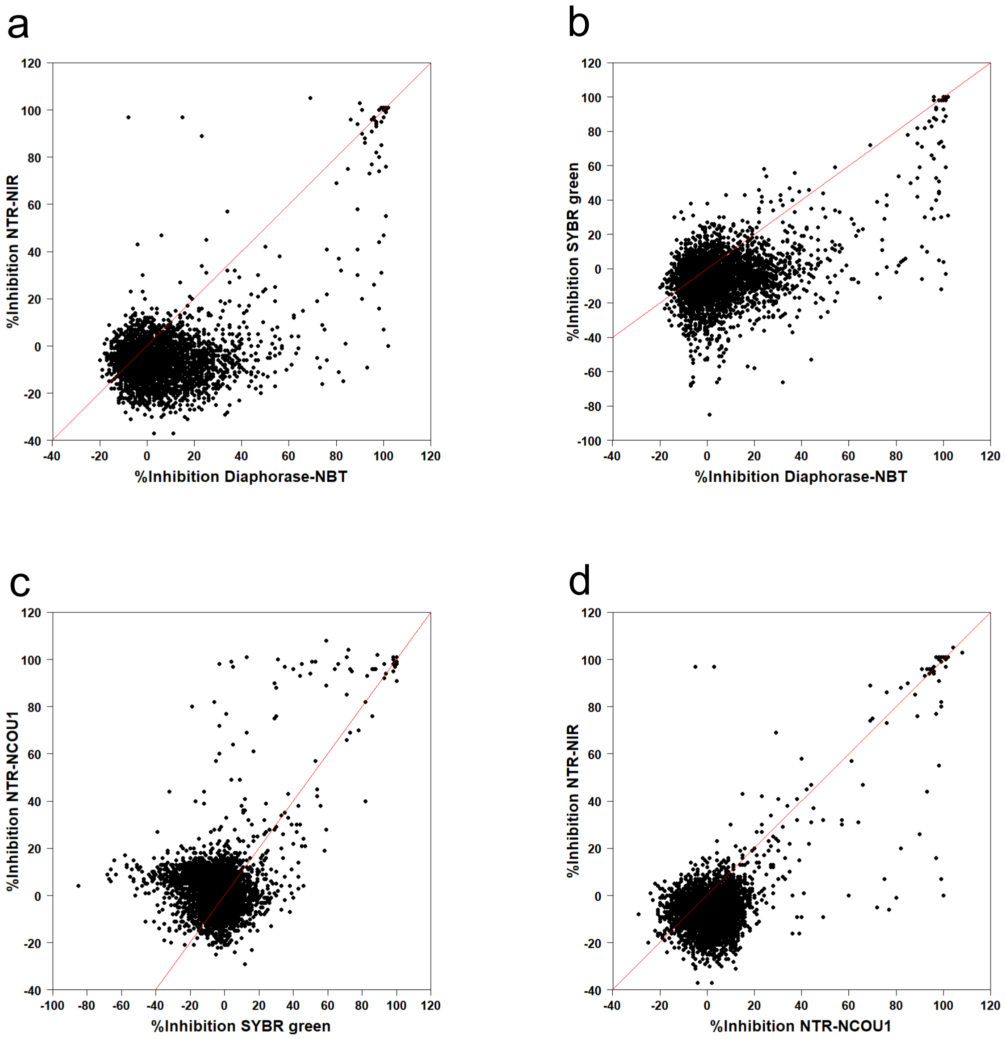


Supplementary Figure 3. Correlations of %Inhibition between assays employing diaphorase-NBT and NTR-NIR (a), diaphorase-NBT and SYBR green (b), SYBR green and NTR-NCOU1 (c), and NTR-NCOU1 and NTR-NIR (d). The red line indicates the line of equity (y = x). NBT: nitroblue tetrazolium, NTR: nitroreductase, NCOU1: nitrobenzyl-umbelliferone 1, NIR: near-infrared.

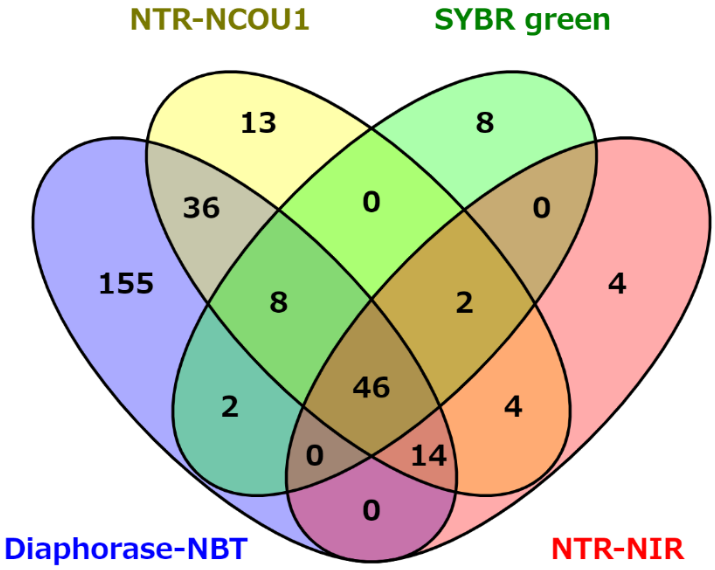


Supplementary Figure 4. Venn diagram of the hit compounds identified by each assay method. NBT: nitroblue tetrazolium, NTR: nitroreductase, NCOU1: nitrobenzyl-umbelliferone 1, NIR: near-infrared.

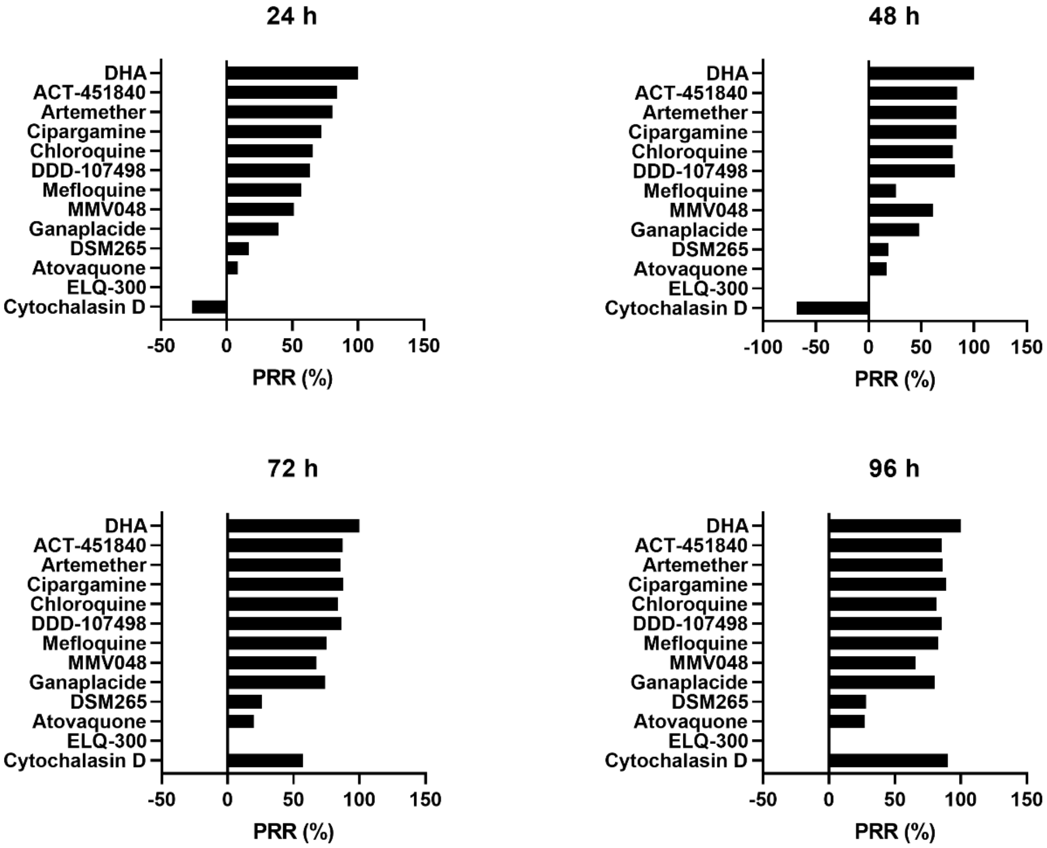


Supplementary Figure 5. HT-PRR of test compounds. The number of live parasites following exposure to 25 μM test compounds for 24, 48, 72, and 96 hours was quantified by the PfLDH-NCOU1 assay. Note that the values of PRR (%) were calculated by normalization of signal intensities between 100% [dihydroartemisinin (DHA)] and 0% (ELQ300). Data as presented as the mean (n=2).

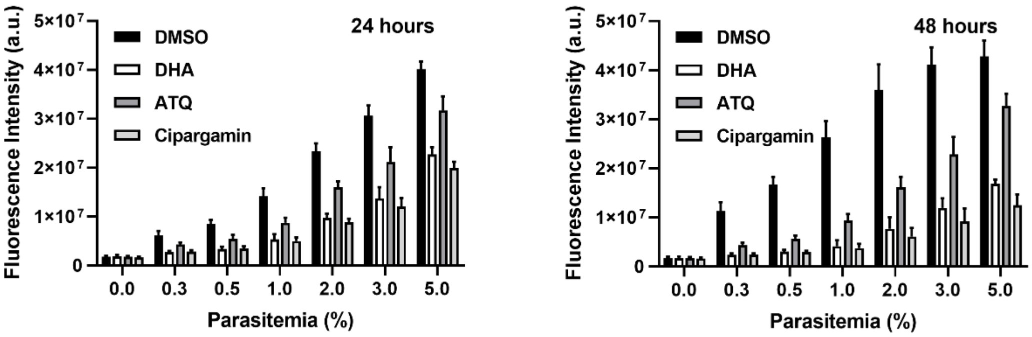


Supplementary Figure 6. Parasite killing by dihydroartemisinin (DHA), atovaquone (ATQ), and cipargamin. Parasites at different parasitemia levels were exposed to compounds for 24 and 48 hours. Data are presented as the mean ± standard deviation (SD) (n=2). DMSO: dimethylsulfoxide, a.u.: arbitrary unit.

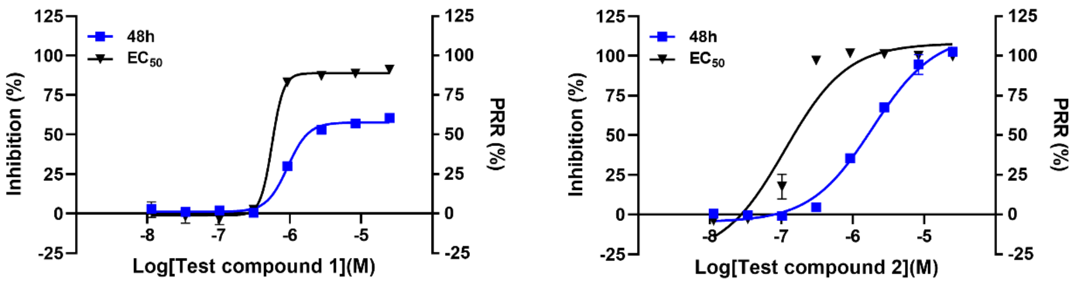


Supplementary Figure 7. Comparison of parasite killing rate for 48 hours and growth inhibition of 2 test compounds. Mixed stage parasites were treated with test compounds for 48 hours parasite reduction ratio (PRR). For growth inhibition assay, 0.3% synchronized ring stage parasites were treated with test compounds for 72 hours. Live parasites of both assays were quantified by PfLDH-NTR assay. Data are presented as the mean ± standard deviation (SD) (n=2). NTR: nitroreductase.
